# A reduction in voluntary physical activity during pregnancy in mice is mediated by prolactin

**DOI:** 10.1101/2020.09.10.292466

**Authors:** S.R. Ladyman, K.M. Carter, Z. Khant Aung, D. R. Grattan

## Abstract

As part of the maternal adaptations to pregnancy, mice show a rapid, profound reduction in voluntary running wheel activity (RWA) as soon as pregnancy is achieved. Here, we evaluate the hypothesis that prolactin, one of the first hormones to change secretion pattern following mating, is involved in driving this suppression of physical activity levels during pregnancy. We show that prolactin can acutely suppress RWA in virgin female mice, and that conditional deletion of prolactin receptors (Prlr) from either all forebrain neurons or from GABA neurons prevented the early pregnancy-induced suppression of RWA. Deletion of Prlr specifically from the medial preoptic area, a brain region associated with multiple homeostatic and behavioural roles including parental behaviour, completely abolished the early pregnancy-induced suppression of RWA. Our data demonstrate a key role for prolactin in suppressing voluntary physical activity during early pregnancy, highlighting a novel biological basis for reduced physical activity in pregnancy.

## Introduction

Pregnancy and lactation represent profound physiological challenges. The sustained changes in metabolic rate typical of pregnancy have been described as “at the limits of human physical capability”, similar to that expended in extreme physical activity such as an ultramarathon, but over a longer timeframe (1). To enable the pregnant female to cope with these demands, hormonal changes associated with pregnancy drive a wide range of adaptations to maternal physiology, and in particular, complex changes in metabolic function (2). Using a mouse model to characterize metabolic adaptations in pregnancy, we have shown that along with increases in energy intake, pregnant females rapidly lower their energy expenditure and physical activity levels, as measured by daily voluntary running wheel activity (3). This profound change in behavior is likely important to offset the prolonged increases in basal metabolic rate characteristic of pregnancy (3–5). Remarkably, however, the reduction in voluntary physical activity starts very early in pregnancy before there is any significant increase in metabolic load, even before implantation (3, 6), and thus, can be considered an anticipatory adaptation in preparation for the future metabolic demands. Based on the very rapid change in behavior, we hypothesized that this reduction in physical activity must be mediated by the hormonal changes associated with the maternal recognition of pregnancy. In rodents, one of the very first changes in hormones in pregnancy is the mating-induced initiation of twice-daily prolactin surges that are required to maintain corpus luteum function in the ovary to sustain the pregnancy (7). Here, we have investigated the role of prolactin acting in the brain to mediate the pregnancy-induced suppression of voluntary physical activity.

## Results

### Reduced physical activity during early pregnancy

The presence of a running wheel enables a robust assessment of an animal’s choice to voluntarily engage in exercise or not. Figure 1A depicts daily running wheel activity (RWA) from a single representative female mouse during different reproductive states, showing the cyclical running patterns characteristic of the estrous cycle with increased running preceding each ovulation; profound reductions in RWA during pregnancy and lactation, apart from a brief increase in RWA the night after birth; and the rapid return to virgin levels of RWA following weaning of pups. To enable experiments both within behavioural phenotyping cages and within their normal home cages, we investigated voluntary RWA during pregnancy independently using two different types of running wheels. Our phenotyping cages (Promethion, Sable Systems International) had a traditional upright wheel, while a saucer/low profile wheel was used in the home cages. The upright wheel potentially takes more effort to run on, whereas the saucer wheel has less resistance and the possibility of ‘coasting’ on the wheel may lead to higher RWA being detected (as suggested by the increased average distance in the saucer group vs the upright wheel group, fig1B P<0.0001) (8–10). Regardless of the different equipment, all mice showed a rapid reduction in RWA early in pregnancy (fig1C, significant effect of time P<0.0001, and fig1D, significant effect of physiological state (virgin vs early pregnancy) P<0.0001, Sidak’s multiple comparisons virgin vs early pregnancy: upright wheel P<0.0001, saucer wheel P<0.0001) demonstrating that this is a robust behavioral change induced as early as the first day of pregnancy. To determine if this pregnancy effect on physical activity was specific to RWA, we also examined general activity levels and energy expenditure in female C57B6/J mice *without access to running wheels*. Mice were housed in Promethion metabolic and behavioural monitoring cages before and during pregnancy (as previously described (3)). We observed that pregnancy significantly influenced both energy expenditure (fig 1E, repeated measures ANOVA P=0.0002) and total daily ambulation (fig 1F, repeated measures ANOVA P=0.003). Both energy expenditure (fig 1E, paired t-test P=0.03) and ambulation (fig 1F paired t-test P=0.03) were significantly reduced in early pregnancy compared to the virgin state. This suggests that early pregnancy is associated with reduced physical activity, independent of the actual form of physical activity measured.

**Figure 1:**
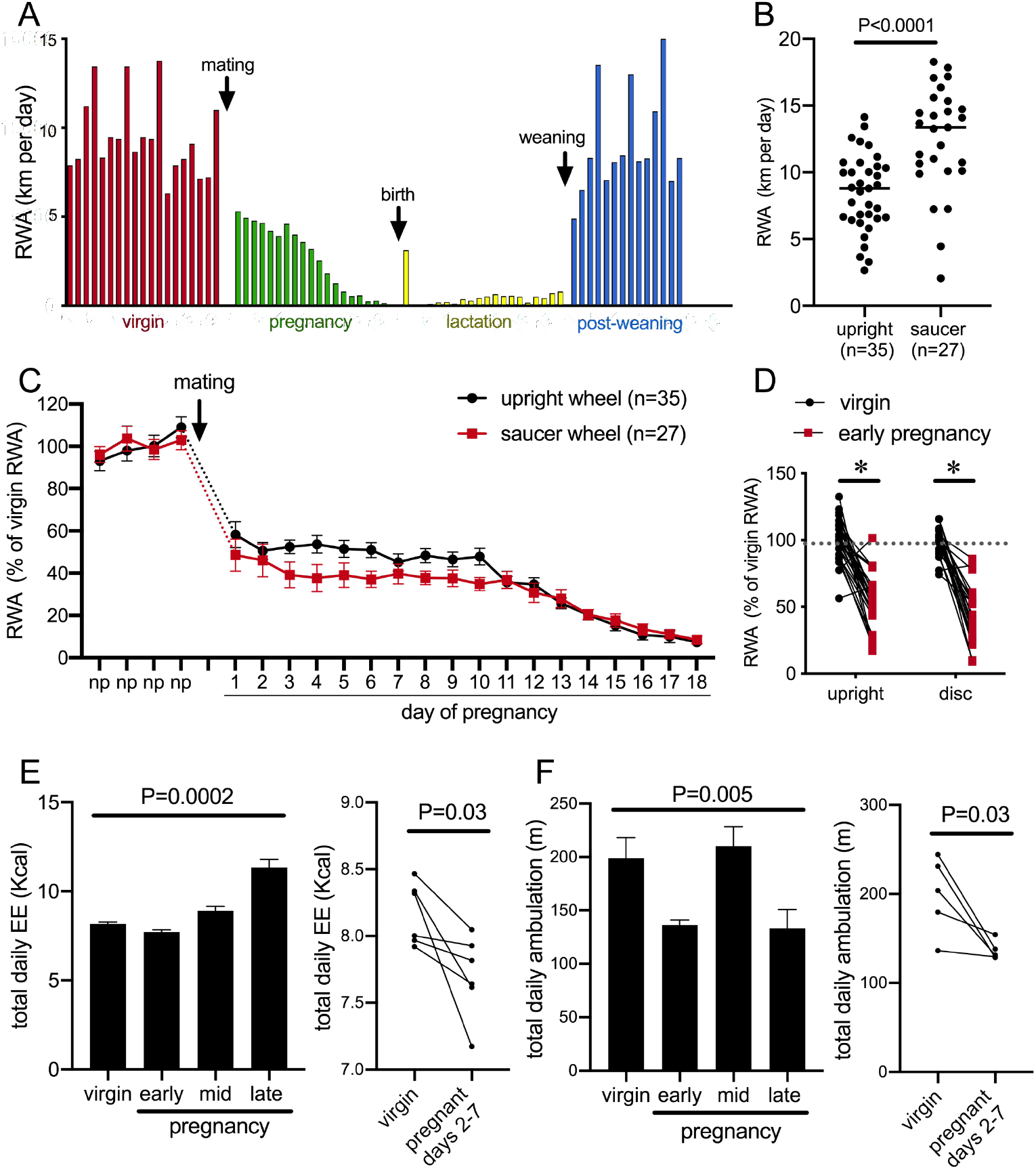
Pregnancy rapidly decreases voluntary physical activity including running wheel activity (RWA). **A**: daily RWA of one representative mouse through one cycle of reproduction (virgin state, pregnancy, lactation and post weaning of her pups). Each bar represents RWA on one day. **B**: RWA in virgin female mice with access to a traditional upright wheel or a low-profile saucer wheel. **C**: RWA in female mice, with either traditional upright wheel or a low-profile saucer wheel, before and after successful mating. RWA activity for each mouse is expressed as a percentage of their average daily levels in the virgin state. **D**: Daily RWA for the first three days of pregnancy expressed as a percentage of each mouse’s RWA level in the virgin state. Dotted line indicates virgin level, and * indicates a significant difference between virgin RWA and early pregnancy RWA (Effect of physiological state P<0.0001, *post hoc* virgin vs early pregnancy upright wheel: P<0.0001, saucer wheel: P<0.0001), n values same as fig 1C. **E**: (*Left*) Bars represent total daily energy expenditure (EE) in the virgin state and different time points during pregnancy (early = days 2-7, mid = days 8-13, late = days 14-18) (repeated measures one way ANOVA, n=6). (*Right*) Points and lines represent the change in total daily EE for each individual mouse between the virgin state and early pregnancy (days 2-7)(t-test, n=6). **F**: (*Left*) Bars represent total daily ambulation in the virgin state and different time points during pregnancy (early = days 2-7, mid = days 8-13, late = days 14-18) (repeated measures one way ANOVA, n=5). (*Right*) Points and lines represent the change in total daily ambulation for each individual mouse between the virgin state and early pregnancy (days 2-7) (t-test, n=5).

### Acute effects of prolactin on physical activity and behavior

The immediate change in behavior following the onset of pregnancy suggested that it was induced by hormonal changes very early in pregnancy. We hypothesized that prolactin was the most likely candidate, due to the rapid induction of twice daily prolactin surges induced by mating in rodents. This idea was supported by the observation that the reduction in RWA persisted throughout pregnancy and lactation (Fig 1A), conditions characterized by high prolactin (7). To determine if prolactin can acutely influence physical activity, we investigated the effects of acute prolactin treatment on physical activity in non-pregnant mice. In addition to RWA in their home cages, a number of other indices of physical activity were assessed in various behavioural tests including both novel and familiar environments. C57BL/6J female mice (metestrous phase of the estrous cycle) that had been housed with a running wheel for at least 3 weeks, were injected i.p. with either prolactin (5mg/kg) or vehicle 30 minutes before the start of the dark phase of the light/dark cycle and running wheel activity was monitored. On the following metestrus, mice were injected with the alternative treatment (prolactin or vehicle) such that all mice received both treatment and control in a random order. Prolactin treatment lead to a significant reduction in RWA (fig 2A, repeated measures two-way ANOVA, interaction time x treatment P<0.0001 and fig 2B, paired t-test P=0.007), particularly in the first three hours of the dark phase (fig 2A, Sidak’s multiple comparisons test: Vehicle vs Prolactin 1h P=0.0009, 2h P=0.0173, 3h P=0.0248) when female mice normally engage in their maximal levels of RWA (8, 10). No effect of acute prolactin treatment on RWA activity was observed in male mice (fig 2C and D).

**Figure 2:**
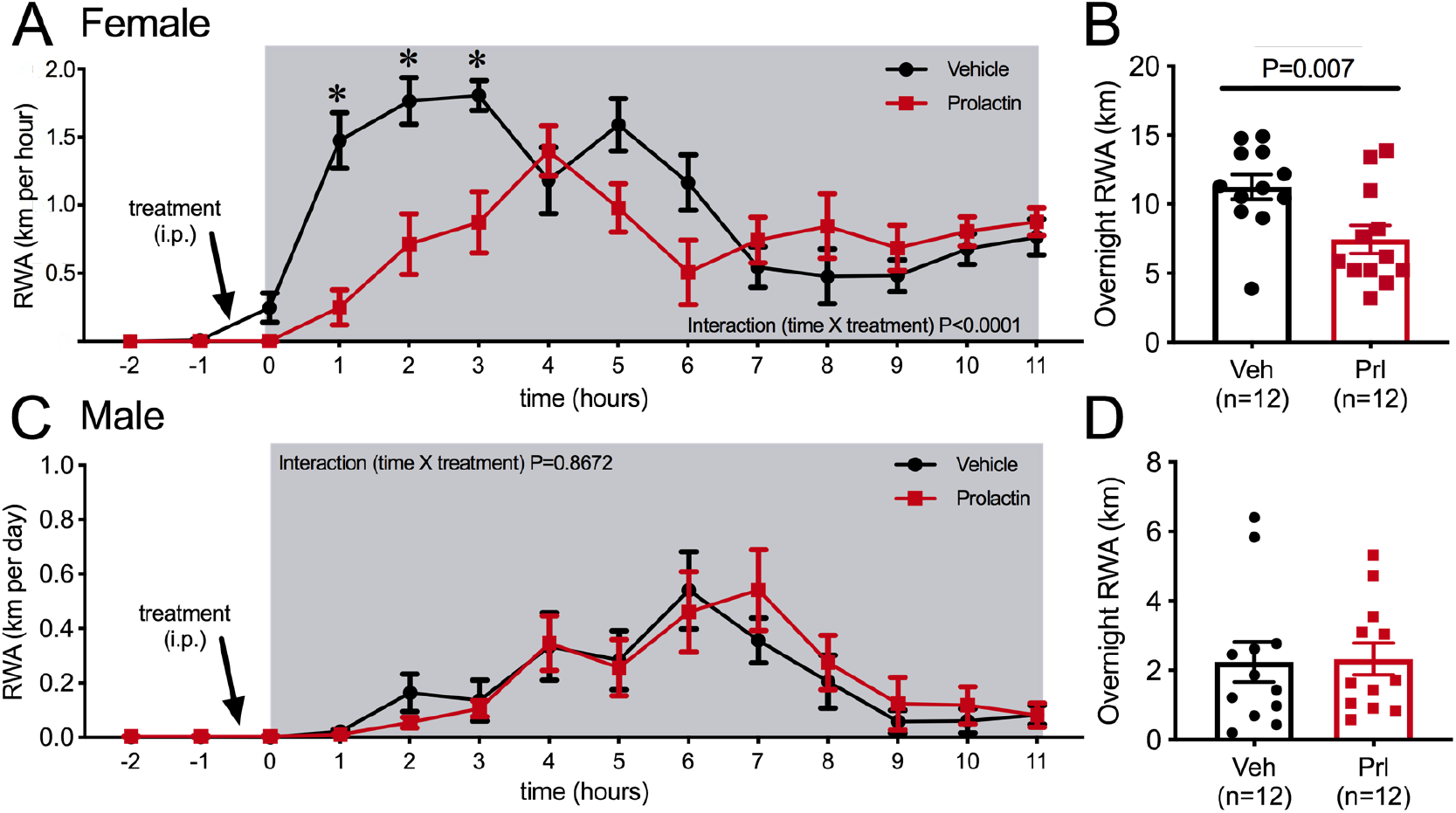
Acute effects of exogenous prolactin on physical activity. **A**: Acute prolactin treatment significantly reduced running wheel activity (RWA) in female mice (**A** and **B**) but not in male mice (**C** and **D**). **A**, **C**: Lines indicate running wheel activity per hour in virgin mice (metestrous phase of estrous cycle for female mice) treated with either vehicle or prolactin (5mg/kg i.p.) 30 minutes before the start of the dark phase (dark phase indicated by grey shading). All mice (n=12 per sex) received both treatments in a randomized order. * indicates time points that showed a significant difference (*Post hoc* analysis P<0.05). **B, D**: Bars represent total 12 hour RWA following either vehicle or prolactin (5mg/kg i.p.) as described above.

In a novel environment, such as in an open field or the elevated plus maze paradigm, prolactin injection an hour prior to testing did not influence distance travelled by female mice (fig 3A and B), suggesting there is not a generalized effect on locomotion. To enable more detail investigation of physical activity levels within the familiar home-cage environment, ambulatory movement was next examined in C57BL/6J female mice housed in Promethion metabolic phenotyping cages, without access to running wheels. Mice were habituated to these cages along with handing and removal from cages for at least a week. On the metestrous phase of the estrous cycle, mice received an i.p. injection of either prolactin (5mg/kg) or vehicle 30 minutes before the start of the dark phase of the light cycle, in a counter balanced manner as described above. Prolactin treatment lead to a significant reduction in ambulation during the 12 hours of the dark phase of the light cycle (fig 3C, repeated measures ANOVA Interaction time x treatment P=0.016), although this was a subtle effect and only evident in the late stages of the dark phase (fig 3C, Sidak’s multiple comparisons test: vehicle vs prolactin 12h P=0.03). The measure of “ambulation distance” in these cages is generated by an algorithm that interprets multiple infra-red beam breaks in a consecutive direction as forward movement or ‘ambulation’. Interestingly, when total counts of XY beam breaks were examined, prolactin treatment tended to increase, rather than decrease, total beam breaks although there was no significant difference detected (fig 3D, repeated measures ANOVA Interaction time x treatment P=0.078). Further examination indicated that prolactin treatment significantly increased fine movement beam breaks, defined as beam breaks that are not in a consecutive direction thus indicating non-ambulatory movement (fig 3E, repeated measures ANOVA Interaction time x treatment P<0.0001, *post hoc* analysis: vehicle vs prolactin P<0.05 for hours 5 to 12). This increase in fine movement counts was observed both in the first 4 hours (fig 3F, paired t-test P=0.0009) and the total 12 hours (paired t-test P=0.028, data not shown) of the dark phase. To determine what fine movement activities might be influenced by prolactin, another cohort of C57BL/6J female mice were video recorded in their home cages for 40 minutes following prolactin or vehicle injection (as described above). Recordings were analyzed to determine the time spent in various activities, such as nest building and grooming. There was no difference identified in the time spent doing any of these behaviors following prolactin or vehicle injection except for time spent ‘still, off nest’ which was slightly yet significantly reduced in the prolactin treated group (fig 3G, paired student t-tests, ‘still, off nest’: P=0.047). Overall, this set of experiments demonstrated that prolactin reduces voluntary RWA in female mice, but had little or no significant impact on general ambulation levels in either a novel environment or in the home cage. Therefore, while rapid changes in RWA activity in early pregnancy may be driven by mating-induced prolactin surges, there is no evidence to support a similar mechanism underlying reductions in ambulation in early pregnancy (fig 1F).

**Figure 3:**
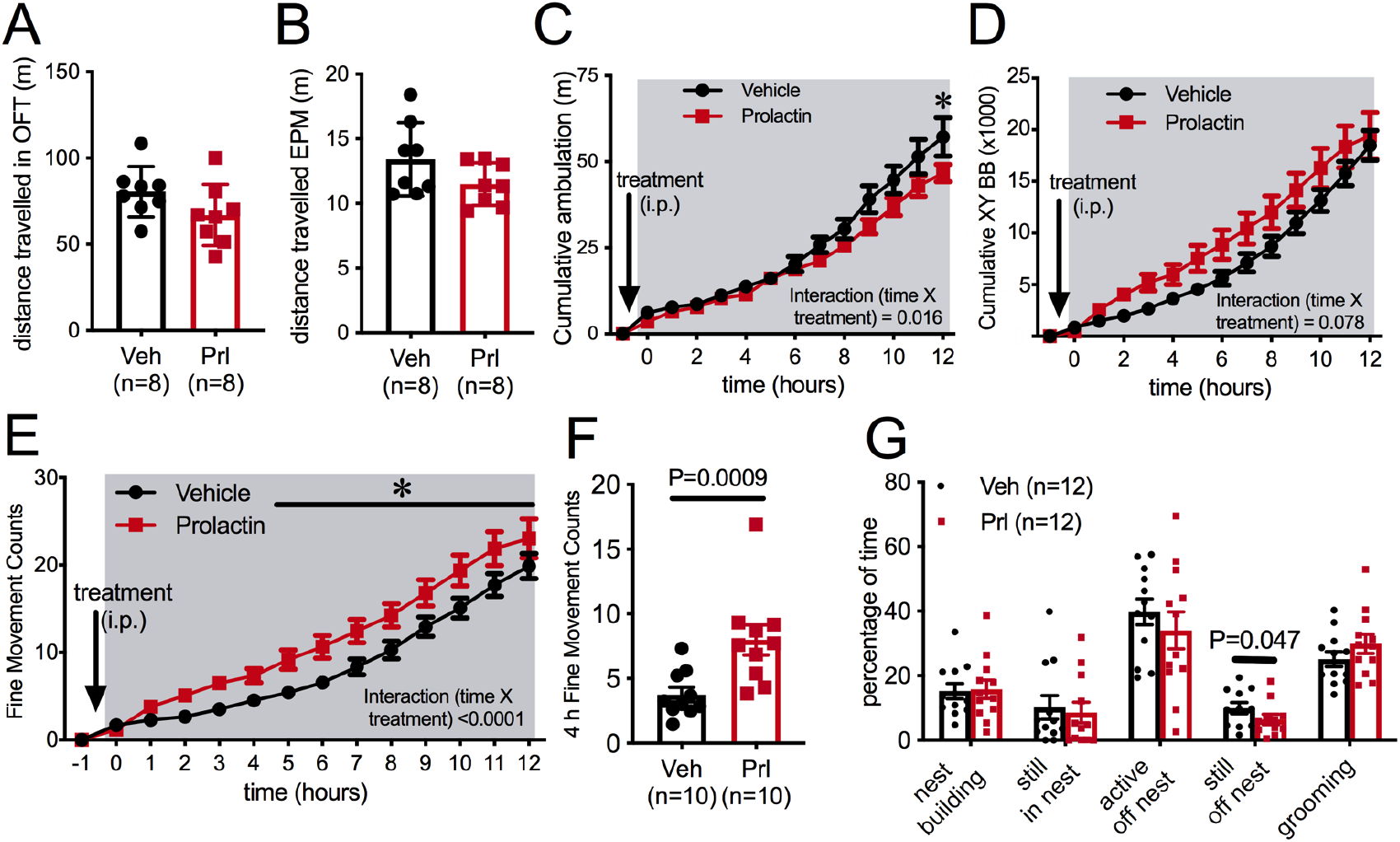
Prolactin does not acutely decrease locomotion. Bars represent total distanced travelled in an open field test (OFT) of 30 minute duration (**A**) or elevated plus maze (EPM) of 5 minute duration (**B**) carried out in the dark phase of the light cycle under dark conditions following either vehicle or prolactin (5mg/kg i.p.) treatment in virgin female mice mice (n=8 per group, metestrous phase of estrous cycle)(t-test). **C-F**: Acute prolactin had a subtle effect on ambulation in home cage. Lines represent cumulative ambulation (**C**), total X + Y beambreaks (**D**) and fine movement (activity that is not ambulation)(**E**) of virgin female mice in the metestrous phase of the estrous cycle in home cage during the dark phase of the light cycle after vehicle or prolactin (5mg/kg, i.p.) treatment (Repeated measures ANOVA). **F**: Bars represent total fine movement counts in the first four hours following the start of the dark phase (paired t-test). **G**: Acute prolactin did greatly impact on time mice spent engaging in different behaviours in their home cages (Paired t-test: still off nest Vehicle vs Prolactin treatment P=0.047 (n=11 due to exclusion of data due to outlier (Rout outlier test, outlier much higher value than the rest of the group)), all other behaviours P>0.05). Bars represent percentage of time spend mice were engaged in various activities in their home cages (total test time 40 minutes) following vehicle or prolactin (5mg/kg, i.p.) treatment (n = 12 mice, all mice received both vehicle and prolactin in a randomized order, paired t-test). * indicates a significant difference, based on the indicated statistical test.

### Prolactin receptors in forebrain neurons are necessary for pregnancy-induced suppression of running wheel activity

To test the hypothesis that prolactin is acting in the brain to mediate the pregnancy-induced suppression of RWA, we measured RWA in a mouse line lacking Prlr in forebrain neurons (Prlr^lox/lox^ mice crossed with CamKinase2-Cre mice (Prlr^lox/lox^/CamK-Cre), as previously described (11)) (see fig 4 Suppl. 1A for validation of Prlr deletion in these mice). RWA was monitored for at least two weeks in the virgin state, then mice were mated and RWA was monitored throughout pregnancy, along with body weight and food intake. These mice do not show normal 4-5 day estrous cycles (fig 4 Suppl. 1B) due to hyperprolactinemia caused by a lack of negative feedback in the hypothalamus (11). The high prolactin acts in the ovary to prolong progesterone secretion after each ovulation, resulting in serial pseudopregnancy-like phases of around 10-12 days between ovulations. We observed a trend for reduced RWA in virgin Prlr^lox/lox^/CamK-Cre compared to control mice (Prlr^lox/lox^) (fig 4A, t-test P=0.058), and in particular, the absence of the cyclical pattern of elevated RWA prior to ovulation that has previously been described (3, 12-14) (fig 4B). Despite the abnormal cycles and slightly increased body weight in the virgin state (fig 4 Suppl 1B and C), these mice are able to become pregnant and maintain a pregnancy, showing no differences in fetus number and uterus mass in late pregnancy or litter size on day 4 of lactation (fig 4 Suppl. 1D-H).

**Figure 4:**
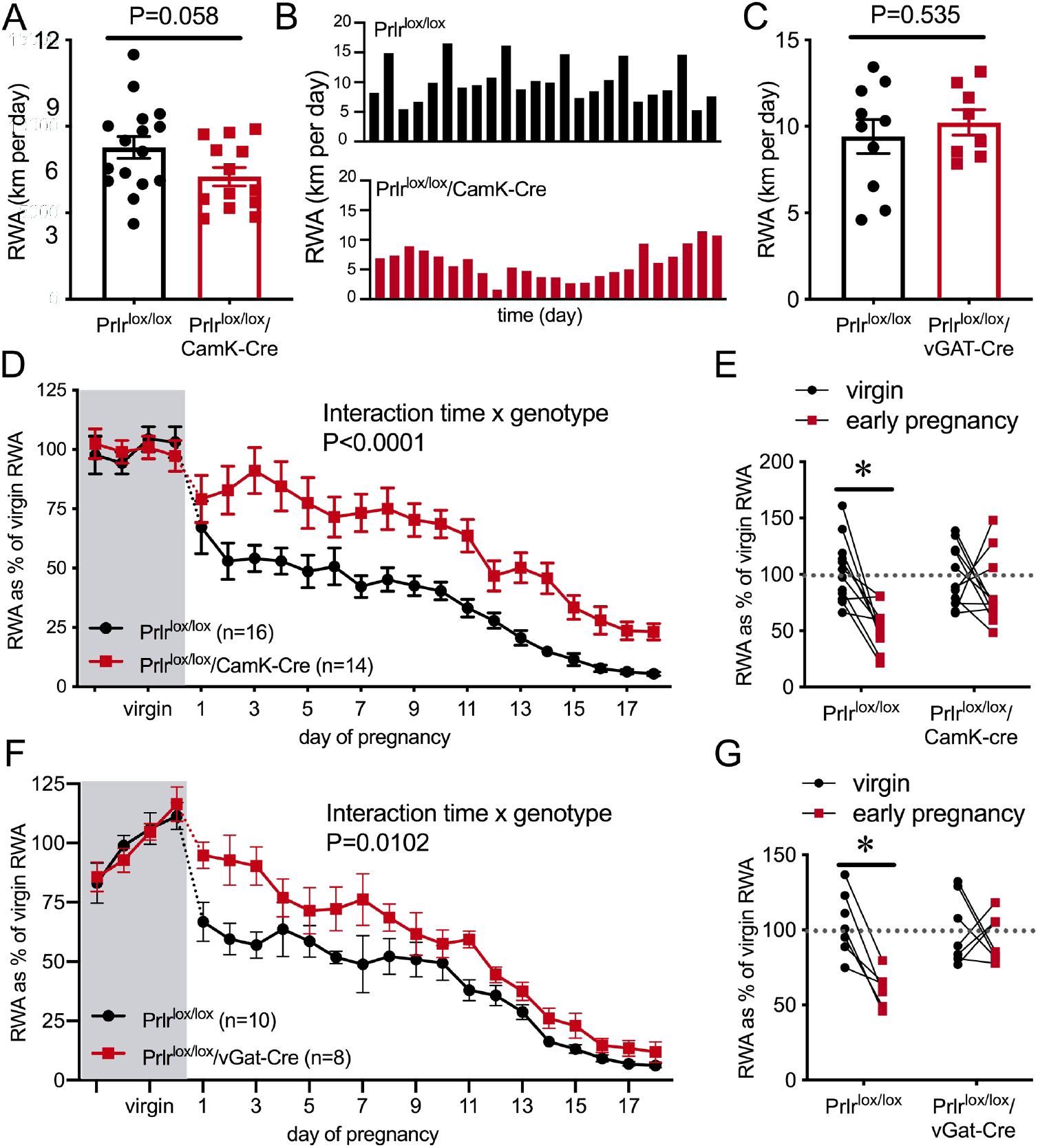
Pregnancy-induced suppression of running wheel activity (RWA) is attenuated in mice lacking Prlr in forebrain neurons. **A, C**: Bars represent average daily RWA in control (Prlr^lox/lox^) and mice lacking Prlr in forebrain neurons (**A**)(Prlr^lox/lox^/CamK-Cre n=14, Prlr^lox/lox^, n=16) or GABA neurons (**C**) (Prlr^lox/lox^/vGAT-Cre n=8, Prlr^lox/lox^, n=10)(t-test). **B**: Representative examples of daily RWA in one control and one Prlr^lox/lox^/CamK-Cre mouse demonstrating a cyclical pattern of increased RWA every 4^th^ day in control mice whereas this pattern is not clearly observed in mice with a deletion of Prlr in forebrain neurons. Each bar represents RWA on a single day. **D, F**: Forebrain Prlr are required for pregnancy-induced suppression of running wheel activity. Line represents RWA before and after successful mating in control mice (Prlr^lox/lox^) and mice lacking prolactin receptors in forebrain neurons (**D**) (Prlr^lox/lox^/CamK-Cre n=14, Prlr^lox/lox^, n=16, total daily RWA per animal shown in Figure 4 – source data 1) or mice lacking prolactin receptors in GABA neurons (**F**) (Prlr^lox/lox^/vGAT-Cre n=8,Prlr^lox/lox^, n=10, total daily RWA per animal shown in Figure 4 – source data 2) mice. RWA activity for each mouse is expressed as a percentage of their individual average daily levels in the virgin state and analysis by mixed effects model. **E, G**: Bars represent daily RWA for the first three days of pregnancy expressed as a percentage of each mouse’s RWA level in the virgin state. Dotted line indicates virgin level, and * indicates a significant difference between virgin RWA and early pregnancy RWA (**E** Forebrain deletion of Prlr: *post hoc* virgin vs early pregnancy: Prlr^lox/lox^: P=0.0002, Prlr^lox/lox^/CamK-Cre: P=0.3899, **G** GABA neuron deletion of Prlr: *post hoc* virgin vs early pregnancy: Prlr^lox/lox^: P=0.0007, Prlr^lox/lox^/vGat-Cre: P=0.7888).

The Prlr^lox/lox^/CamK-Cre mice failed to show the characteristic full decrease in running in early pregnancy, and in fact showed significantly increased running wheel activity compared to control mice throughout pregnancy (fig 4D, interaction genotype x time P<0.0001). Even when heavily pregnant on days 16-18 of pregnancy, the Prlr^lox/lox^/CamK-Cre animals continued to run more on the wheel that controls. When analyzing the overall change in RWA across the first week of pregnancy, Prlr^lox/lox^/CamK-Cre mice did not show a significant reduction in RWA, whereas control mice reduced RWA to approximately 50% of non-pregnant levels in the first few days of pregnancy (fig 4D, interaction genotype x time P=0.0385, *post hoc* for significant effect of time: Prlr^lox/lox^ P=0.002, Prlr^lox/lox^/CamK-Cre P=0.3899). In the presence of a running wheel, Prlr^lox/lox^/CamK-Cre mice gained slightly, yet significantly, less weight by the later stages of pregnancy (fig 4 Supplementary 1H) however they did not show any change in daily food intake (fig 4 Suppl. 1G). To eliminate the possibility that the loss of Prlr in the brain might cause exaggerated stress responses in the pregnant females (15, 16) that could influence acute RWA (17), we examined anxiety-like behavior in late pregnant (day 15/16) Prlr^lox/lox^/CamK-Cre and control mice using the elevated plus maze, as described previously (18). Time spent in the open arms of the EPM was similar in both groups, suggesting no difference in anxiety-like behavior in the pregnant Prlr^lox/lox^/CamK-Cre and controls (fig 4 Supplementary 1I).

While the data showing a lack of rapid pregnancy-induced suppression of RWA in Prlr^lox/lox^/CamK-Cre were clear, the elevated prolactin (11) and abnormal estrous cycles (fig 4 Suppl. 1B) were a potential confound in this mouse line. Hence, we aimed to replicate these experiments using a mouse line with a specific deletion of Prlr from GABA neurons (Prlr^lox/lox^ crossed with vGAT-cre mice, as previously described (11)) (fig 4 Suppl 2A). These mice have more limited, although still extensive, deletion of prolactin receptors throughout the brain (11, 19). Importantly, however, Prlr are not deleted from the majority of hypothalamic dopamine neurons, meaning that prolactin levels are normal and they have normal 4-5 day estrous cycles (11). Prlr^lox/lox^/vGat-cre mice have similar body weight in the virgin state as controls (fig 4 Suppl. 2B), can carry a pregnancy to term, and both genotypes had similar sized litters (fig 4 Suppl. 2C). Virgin Prlr^lox/lox^/vGat-cre and control (Prlr^lox/lox^) mice had similar levels of RWA (fig 4C). In contrast, however, Prlr^lox/lox^/vGat-cre mice had markedly higher RWA throughout pregnancy compared to controls (fig 4F, Interaction time x genotype P=0.010). Consistent with observations in Prlr^lox/lox^/CamK-Cre mice, Prlr^lox/lox^/vGat-Cre mice did not show the rapid reduction in RWA in early pregnancy, whereas control mice showed the characteristic pregnancy-induced immediate reduction in RWA (fig 4G, Interaction genotype x time P=0.0162, *post hoc* for significant effect of time: Prlr^lox/lox^ P=0.0007, Prlr^lox/lox^/vGat-Cre P=0.7888). Again, similar to Prlr^lox/lox^/CamK-Cre, no differences in anxiety-like behavior were detected in pregnant Prlr^lox/lox^/vGat-Cre mice (fig 4 Supplementary 2D). There was no difference in body weight gain or food intake during pregnancy in Prlr^lox/lox^/vGat-Cre mice compared to controls (fig 4 Supplementary 2E and F). Overall, the attenuated reduction in RWA in early pregnancy in both Prlr^lox/lox^/CamK-Cre and Prlr^lox/lox^/vGat-Cre strongly indicates a role for prolactin action in the brain, particularly through a population of GABAergic neurons, driving early changes in RWA as soon as pregnancy is initiated.

### Identifying the neuronal populations involved in prolactin-induced suppression of RWA during pregnancy

Based on a report that ablation of arcuate nucleus kisspeptin neurons markedly suppressed RWA in female mice (20), we next examined the role of these neurons in mediating the pregnancy-induced decrease in RWA. We have shown that arcuate kisspeptin neurons express Prlr (21, 22) and that prolactin is likely to inhibit their activity (23), although it should be noted that only a small proportion of these kisspeptin cells are GABAergic (24). Mice with Prlr deleted from kisspeptin neurons were generated by crossing Kiss1-cre mice (25) with Prlr^lox/lox^ mice (Prlr^lox/lox^/Kiss1-cre) (23). We have previously shown that these mice have Prlr deleted specifically in the arcuate nucleus kisspeptin neurons (fig 5A), and that they have normal estrous cycles (23). Overall, there was a significant effect of pregnancy on RWA activity in both Prlr^lox/lox^/Kiss1-cre and control Prlr^lox/lox^ mice (fig 5B, effect of pregnancy P<0.0001) but no difference between the genotypes. On closer analysis, the effect of pregnancy on RWA in the first few days of pregnancy (days 1-3) was subtly reduced in Prlr^lox/lox^/Kiss1-cre mice compared with control mice (fig 5C, interaction genotype X early pregnancy P=0.01, *post hoc* for significant effect of genotype: early pregnancy P=0.0162), but not to the same extent seen in the CamK- or GABA-specific Prlr knockout mice. Thus, these data suggest the possibility that prolactin action on arcuate kisspeptin neurons may play a small contributing role in the initial reduction in RWA during pregnancy, but are not major factors in mediating this response.

**Figure 5:**
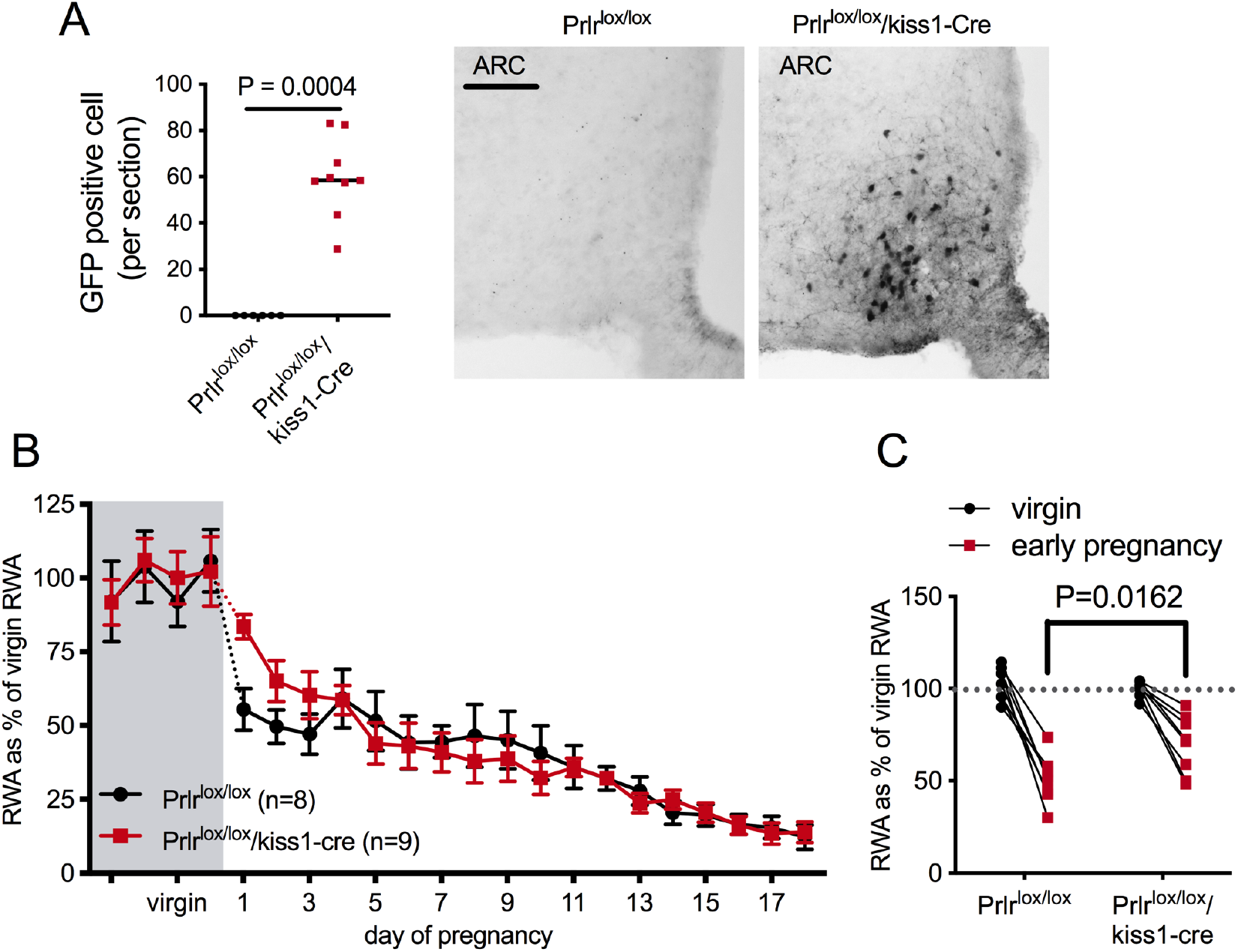
Effect of Prlr deletion from arcuate nucleus kisspeptin neurons on running wheel activity during pregnancy. **A:**Bars represent number of GFP positive cells (black staining), indicative of Prlr gene deletion, in the arcuate nucleus of Prlr^lox/lox^/kiss1-Cre female mice alongside representative images, n=6-9 per group (Mann Whitney test) **B**: Mice with Prlr deleted from arcuate nucleus kisspeptin neurons show a subtle difference in early pregnancy-induces changes in RWA. Line represents RWA before and after successful mating in mice lacking Prlr in arcuate nucleus kisspeptin neurons (Prlr^lox/lox^/kiss1-Cre n=9) and control (Prlr^lox/lox^, n=8) mice. RWA activity for each mouse is expressed as a percentage of their average daily levels in the virgin state, total daily RWA per animal shown in Figure 5 – source data 1. **C**: Early pregnancy-induced decrease in RWA is reduced in mice lacking Prlr in arcuate nucleus kisspeptin neurons. Bars represent daily RWA for the first three days of pregnancy expressed as a percentage of each mouse’s RWA level in the virgin state. Dotted line indicates average virgin level. Both Prlr^lox/lox^ and Prlr^lox/lox^/kiss1-Cre mice showed a significant reduction in RWA following successful mating (*post hoc* analysis virgin vs early pregnancy: Prlr^lox/lox^ P<<0.0001, Prlr^lox/lox^/kiss1-Cre P<0.0001), however this decrease in RWA in early pregnancy was not as low in the Prlr^lox/lox^/kiss1-Cre as that seen in the controls (*post hoc* analysis Prlr^lox/lox^ vs Prlr^lox/lox^/kiss1-Cre: early pregnancy P=0.0162).

RWA is inherently rewarding for rodents (reviewed in (26)), and we hypothesized prolactin might alter functions of the midbrain dopamine-mediated reward pathways to influence RWA. While we do not see Prlr expression in the cell bodies of these dopamine neurons in the ventral tegmental area (VTA) (27), there are Prlr-receptor positive fibers in this region, and we have characterized a projection from Prlr-expressing neurons in the medial preoptic area (MPOA) to the VTA (19). Moreover, approximately 50% of Prlr-expressing neurons in the MPOA are GABAergic (19). Hence, our final hypothesis was that Prlr action in the MPOA may be a key site to integrate prolactin action with the reward pathway that promotes RWA. To investigate this possibility, we used an adeno-associated virus (AVV) to deliver Cre recombinase specifically into the MPOA of adult female Prlr^lox/lox^ mice, with controls consisting of AAV-mCherry delivered to Prlr^lox/lox^ mice. Following recovery from stereotaxic surgery for virus delivery, mice were housed with a running wheel for at least three weeks, then mated and RWA was monitored throughout pregnancy. The Prlr^lox/lox^ construct is designed such that Cre-mediated recombination will activate GFP expression and thus report successful Prlr deletion (fig 6A). Similar to what we have reported previously, AAV-cre injection into the MPOA of Prlr^lox/lox^ mice removed functional prolactin responses from this region, as determined by prolactin-induced pSTAT5 (fig 6A) and led to the abandonment of pups following birth (fig 6B), confirming the role of Prlr in this area for normal maternal behavior in lactation (19). Mice with a specific deletion of Prlr in the MPOA did not show the immediate change in RWA once pregnant and maintained higher levels of RWA throughout pregnancy compared to control injected Prlr^lox/lox^ mice (fig 6C, interaction virus injection X day of pregnancy P<0.0001). Strikingly, mice lacking Prlr in the MPOA did not demonstrate any reduction in RWA in the first 3 days of pregnancy (fig 6C, interaction virus injection X early pregnancy P=0.0267, *post hoc* analysis for effect of early pregnancy: control injection P=0.01, AAV-Cre injection P=0.9687), a time when dramatic reductions are occurring in controls. These data indicate that the MPOA is the key site for prolactin-induced suppression of RWA during early pregnancy. Overall, our data would suggest that a population of prolactin-responsive GABAergic neurons in the MPOA induces this profound behavioral change in early pregnancy.

**Figure 6:**
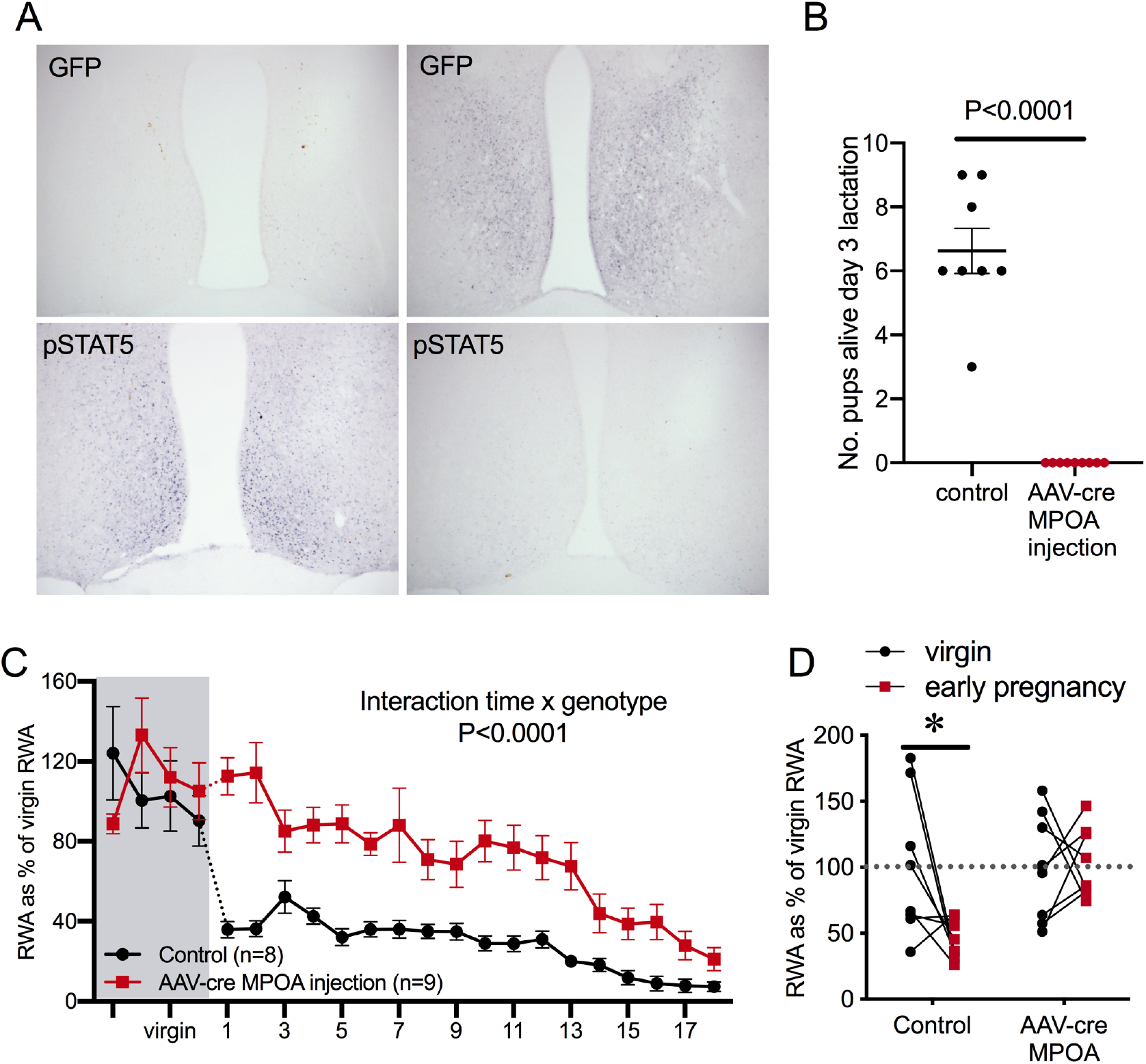
MPOA prolactin sensitive neurons mediate effect of prolactin on running wheel activity (RWA) during pregnancy. **A**: Representative images of coronal sections through the MPOA showing immunohistochemical staining for extensive green fluorescence protein (GFP) and absence of pSTAT5 in AAV-cre injection Prlr^lox/lox^ mice (right) and absence of GFP and extensive pSTAT5 in control virus injection or AAV-cre injection Prlr^lox/lox^ mice. **B**: No pups from MPOA AAV-cre injection Prlr^lox/lox^ mice survived to day 3 of lactation (Mann Whitney test). **C**: Prlr in the MPOA is required for pregnancy-induced suppression of running wheel activity. Line represents RWA before and after successful mating in mice lacking Prlr in the MPOA following AAV-cre injection (AAV-cre MPOA injection, n=9) and control (control virus injection, n=8) mice. RWA activity for each mouse is expressed as a percentage of their average daily levels in the virgin state (Mixed effect model), total daily RWA per animal shown in Figure 6 – source data 1. **D**: Pregnancy-induced decrease in RWA is attenuated in mice lacking Prlr in MPOA. Bars represent daily RWA for the RWA and early pregnancy RWA (*post hoc* virgin vs early pregnancy: control: P=0.01 as indicated by *, AAV-cre MPOA injection: P=0.9687).

## Discussion

In this study, we have shown that the mating-induced release of prolactin, acting through central Prlr, induces a marked reduction in running wheel activity in mice very early in their pregnancy. It seems likely that this profound behavioural response is an important addition to a range of hormone-induced metabolic adaptations that have been characterized to facilitate a positive energy balance during pregnancy, including leptin resistance, increased appetite and reduced energy expenditure (3, 28). From an evolutionary perspective, these can be viewed as adaptive strategies, enabling pregnant females to build up energy reserves in preparation for the pronounced metabolic demands associated with fetal growth and the subsequent lactation (29, 30). In our modern obesogenic environment, however, such changes have become maladaptive, contributing to excessive weight gain during pregnancy with the associated increased risk in pregnancy complications (31). Increasing numbers of women are coming into pregnancy already overweight or obese, and over 50% of women now exceed the Institute of Medicine (IOM) guidelines for optimal weight gain during pregnancy (32–34). Indeed, obesity during pregnancy has been described as the “*most common clinical risk factor encountered in obstetric practice*” (35). As it is difficult to alter energy intake, increasing energy expenditure through exercise during pregnancy has become highly advocated as a therapeutic intervention that produces healthier gestational weight gain (36). However, despite pregnant women being well-informed about the benefits and safety of exercise during pregnancy (37), 60-80% of pregnant women do not engage in physical activity as recommended (38–40). This suggests a disconnect between knowledge and action (37), and poses the question of whether there is a biological bases for decreased physical exercise during pregnancy. Indeed, women report fatigue, pregnancy symptoms and lack of motivation all as barriers to exercise during pregnancy (37). Here, we describe a previously unknown hormonal mechanism that may be contributing to the loss of motivation for exercise in pregnancy.

Our data implicate a population of neurons in the MPOA, highly likely to be GABAergic, in mediating this profound change in behavior. The MPOA is involved in regulating a wide range of homeostatic and behavioural functions, including parental behavior (41), and in particular, prolactin action on maternal behavior (42). We have recently shown that prolactin action in this area is critical to the normal expression of maternal behavior in mice (19). Hence, the prolactin-mediated effect on RWA, described here, might be considered part of the complex adaptive role of the MPOA in promoting appropriate parental responses, triggered by the hormonal changes of pregnancy and lactation. The MPOA has previously been linked to RWA, with the increased running in response to estradiol treatment apparently mediated through estrogen receptor alpha (ERα) in the MPOA (43). The neuronal cell types mediating the RWA response to estradiol have not been elucidated, but an estrogen-responsive MPOA circuit projecting to the VTA has been shown to be critical for maternal behavior (44, 45), and at least some of these ERα-expressing neurons in the MPOA are GABAergic (46). It is possible that estradiol and prolactin may be targeting similar populations of neurons to impact this behavior. Given the strong evidence linking VTA dopamine projections with the rewarding aspects of RWA, this remains a strong candidate for mediating the effect of prolactin on RWA.

Prolactin appears to be critical for most of the initial reduction of RWA in early pregnancy, yet as pregnancy progressed, the neuron-specific Prlr knockout mice still showed reduced RWA compared with non-pregnant animals. It is possible that this is simply due to physical constraints, as growth of the fetuses provides an additional burden on the mothers. However, the fact that very low RWA persists throughout lactation points more to a hormonal mechanism being involved, in addition to the inhibition provided by prolactin. A possible candidate is progesterone, which is known to decrease RWA in rodents through opposing the positive influence of estradiol (13). In mice, progesterone begins to be increased around 48 hours following detection of a copulatory plug and then is high throughout pregnancy until around 24 hours before parturition (47, 48). After birth, progesterone secretion is re-initiated in rodents, due to prolactin-induced rescue of the corpus luteum following a postpartum ovulation. While there is evidence from progesterone receptor knock-out rats that progesterone interactions with this receptor are not required for regulating RWA during the estrous cycle (49), it seems likely that the very high levels of progesterone seen during pregnancy are facilitating the prolactin-induced suppression of RWA as pregnancy advances.

Overall, the data provide novel and convincing evidence of a prolactin-mediated adaptive change in voluntary physical exercise during early pregnancy in mice. Whether such a response also occurs in women requires attention. Prolactin, and its closely related placental analogue, placental lactogen are both highly regulated hormones during human pregnancy, among the top 1% of proteins increasing in the blood at this time (50, 51). These changes occur progressively throughout human pregnancy, as opposed to being an immediate mating-induced increase as seen in rodents, but are still likely to be sufficiently elevated in the blood to be having effects in the brain by the end of the first trimester. It seems likely that these elevated levels of lactogenic hormones are contributing to the broad range of adaptations that are occurring during pregnancy (2, 52). As discussed above, however, in the current environment some of these changes may have become maladaptive and potentially compromise healthy behaviors. Such a possibility needs to be seriously considered in providing obstetric advice to pregnant women.

## Materials and Methods

### Animals

Female mice starting at age 8-12 weeks, were housed in a temperature- and lighting-controlled environment (22 ± 1 C, 12 h light:12 h dark, lights on at 0700h) and allowed access to food and water *ad libitum.* When required, mice were handled daily to monitor estrous cyclicity. All experimental protocols were approved by the University of Otago Animal Ethics Committee. Groups of C57BL/6J mice were used for characterization of activity during pregnancy and to investigate the effects of acute prolactin on activity. Prlr^lox/lox^ mice were generated as previously described (11). CamKII-alpha cre (CamK-cre) mice (53), vGat-cre (54)(Jackson lab stock no. 028862) and kiss1-cre (25) were crossed with Prlr^lox/lox^ mice to generate mouse lines with deletion of Prlr in various neuronal populations.

### Prolactin treatment

Exogenous prolactin was either ovine prolactin (Sigma Aldrich) dissolved in saline or ovine prolactin (obtained from Dr. A. F. Parlow, National Hormone and Pituitary Program, National Institute of Diabetes and Digestive and Kidney Diseases, Torrance, CA) dissolved in PBS/130 mm NaCl (pH 8) to inject at a dose of 5mg/kg (i.p.). Vehicle was either saline or PBS/130 mm NaCl (pH 8). For studies with a counter-balanced design, mice underwent both prolactin injection and vehicle injection, with at least one estrous cycle between treatment days (or 4 days for males). For the effect of acute prolactin on RWA and home cage ambulation, injections were given 30 minutes before the onset of the dark period. All injections were given intra-peritoneally (i.p.) and all mice were habituated to handling. All injections in female mice were carried out in the metestrous phase of the estrous cycle.

### Running wheel activity

All mice were housed individually for the assessment of RWA. For assessment of effect of running wheel type on pregnancy-induced changes in RWA mice with access to an upright wheel were housed in Promethion metabolic and behavioral phenotyping cages for approximately 2 weeks, moved to a cage with a stud male until the day a copulatory plug was detected, at which point they were returned to their Promethion metabolic and behavioral phenotyping cages. For comparison between wheel types, mice in the upright wheel group consisted of C57BL/6J mice and control Prlr^lox/lox^ mice. Mice in the saucer wheel group consisted of Prlr^lox/lox^ mice and were housed with wheel access for at least 3 weeks in the virgin state then similarly treated as those in the upright wheel group. For upright wheel group, running wheel data was collected and processed using MetaScreen and Expedata programs. For saucer wheel group, wheel data was collected and processed using Wheel Manager Software (Version 2.03 MED Associates, Inc).

Studies using Prlr^lox/lox^/CamK-cre, Prlr^lox/lox^/vGat-cre and Prlr^lox/lox^/Kiss1-cre mice (and their controls) recorded running activity from upright wheels in metabolic and behavioral monitoring cages (Promethion, Sable Systems International), while studies with Prlr^lox/lox^ mice with AAV-cre injected into the MPOA (and their controls) used wireless low profile ‘saucer or disc type’ running wheels (Med Associates Inc., Vermont, USA). For all studies, mice were habituated to the running wheels in the virgin state (approximately 2 weeks for upright wheels and at least 3 weeks for low profile wheels). For pregnancy studies, mice were removed from their home cages and housed with a stud male (C57BL/6J) mouse until detection of a plug indicating successful mating. Females were then returned to home cages with wheels and were not disturbed till day 3 or 4 of lactation when number of offspring was assessed. Pregnancy-induced changes in RWA was examined in two cohorts of Prlr^lox/lox^/CamK-cre and control mice at different times, data was pooled as results from the first cohort was repeated in the second cohort. For all other transgenic mouse lines (and their littermate controls) the experiment was only performed on one cohort of mice. Data from the virgin and pregnant state were only included in analysis if mice successfully became pregnant and gave birth.

Our data (Fig 1 and (3)) demonstrate that while there is a wide variation in stable, total daily RWA between different mice, pregnancy induces an approximately 50% decrease in RWA as soon pregnancy is achieved independent of whether a mouse is a ‘high’ or ‘low’ runner. Due to the varying range of stable running levels in individual mice, to assess the pregnancy induced change in behaviour we analysed the percentage change in RWA for each individual mouse as opposed to using total distance per day for each mouse (although this data is provided in source data for each transgenic line). Virgin RWA was calculated for the four days prior to mating as a percentage of the average daily RWA over at least a week following running wheel habituation. RWA activity for each day of pregnancy was also calculated in a similar manner, as a percentage of average daily RWA over at least a week.

### Acute effects of prolactin on RWA

Female and male C57Bl/6J mice (n=12 per sex) were housed individually with running wheels for at least three weeks. Estrous cycle was monitored daily in females and treatment (prolactin or vehicle) was injected on the metestrous phase of the estrous cycle. Mice were administered in a randomized, alternate order such that half received 5 mg/kg injections (i.p.) of vehicle or prolactin first and on the next metestrous cycle the same animals received the alternate treatment. Males were first injected with saline or prolactin and four days later received the alternate treatment. Both groups received injections approximately 30 minutes before the onset of the dark phase. This experiment was carried out in one cohort of mice for each sex.

### Acute effects of prolactin on distance travelled in novel environments

All experiments examining acute effects of prolactin on various physical activity measures in female mice was performed in one cohort of mice, although the experiments for different measures used different cohorts of mice. Female C57Bl/6J mice (n=16) were monitored for stage of estrous cycle, and on metestrus they were injected i.p. with either vehicle or prolactin (5mg/kg) approximately 4 hours into the dark phase of the light/dark cycle. One hour later mice were placed in an open field box and recorded using TopScan software (CleverSys, Inc., VA) for the next 30 minutes. OF testing was carried out under sodium lighting to allow video recording but perceived as ‘dark’ to rodents. After completing a full estrous cycle, on the next metestrus mice were similarly injected with vehicle or prolactin (5mg/kg) approximately 4 hours in to the dark phase of the light/dark cycle and then an hour later mice were placed in an elevated plus maze. During the EPM mice were exposed to white light for the duration of the test. The activity of the mouse was recorded using TopScan software and both OF and EPM tests were analyzed for distance traveled during the time period (OF: 30 minutes, EPM: 5 minutes) using TopScan software.

### Acute effects of prolactin on activity in familiar environment

Female C57Bl/6J mice were housed in Promethion cages and underwent two weeks of handling (removal from cages) and estrous cycle monitoring prior to experiment. On metestrus, mice received an injection of prolactin (5mg/kg) or vehicle approximately 30 minutes before the start of the dark phase of the light/dark cycle. On the metestrous of the following estrous cycle, mice received the alternative treatment as described above. Data was collected from Promethion cages using Metascreen software and processed using Expedata software. End points analyzed included total ambulation distance, total infra-red beam breaks and non-ambulatory movements.

### Measure of anxiety-like behaviour during pregnancy

Pregnant Prlr^lox/lox^/CamK-Cre, Prlr^lox/lox^/vGAtT-Cre and Prlr^lox/lox^ mice underwent an elevated plus maze test on day 15 or 16 of pregnancy similar to that described above but with no prior treatment. The activity of the mouse was recorded and analyzed using TopScan software. At completion of the EPM mice were killed and the uterus, including contents, was weighed and fetal number were recorded.

### Stereotaxic injections of AAV

Female Prlr^lox/lox^ mice (8-12 weeks of age) were anesthetized with isoflurane and placed in a stereotaxic apparatus. Mice received bilateral 0.8μl injections of AAV/DJ-CMV-mCherry-iCre (1.8 × 10^13^ Vector Biosystems) for gene deletion group or AAV/DJ-CMV-mCherry (3.7 × 10^13^ Vector Biosystems) for control group into the MPOA (coordinates were 0.07 anterior to Bregma, 0.3 mm lateral to midline, and depth from top of the brain was adjusted by body weight: <22g 4.7mm depth, >22g 4.9 mm). Injections were given at a rate of 80nL/min, and the syringes were left *in situ* for 3 minutes before and 10 minutes following completion of injection. Mice were left to recover for one week, then housed with running wheels for at least 3 weeks prior to mating. Only mice that showed GFP staining in the MPOA, successfully went through pregnancy and abandoned their litters as previously described (19) were included in the MPOA specific deletion of Prlr group. For the control group only mice that underwent surgery for control virus injection, showed no GFP staining in the MPOA and successfully went through pregnancy were included (inclusion/exclusion for the control group was not influenced by successful nurturing of offspring, although no mice in this group abandoned their pups).

### Immunohistochemistry

Prlr^lox/lox^/CamK-cre, Prlr^lox/lox^/vGat-cre and Prlr^lox/lox^/Kiss1-cre mice have been previously characterized (11, 19, 23). To confirm the deletion of Prlr in this study, immunohistochemistry for prolactin-induced pSTAT5 was used for Prlr^lox/lox^/CamK-cre and Prlr^lox/lox^/vGat-cre lines, while immunohistochemistry for GFP, which is indicative of cre-mediated recombination of the Prlr gene in this Prlr^lox/lox^ mouse line, was used for the Prlr^lox/lox^/Kiss1-cre line. Both prolactin-induced pSTAT5 and GFP immunohistochemistry was used to demonstrate deletion of Prlr in MPOA AAV-cre injected Prlr^lox/lox^ mice. For pSTAT5 immunohistochemistry, mice received an i.p. injection of prolactin (5mg/kg) 45 minutes before perfusion. Mice were anesthetized with sodium pentobarbital then perfused transcardially with 4% paraformaldehyde. Brains were removed and processed for immunohistochemistry for GFP or pSTAT5, as previously described (11). Briefly, for pSTAT5 immunohistochemistry coronal brain sections (30 micron) underwent an antigen retrieval procedure consisting of incubation in 0.01 mM Tris (pH 10) at 90 C for 5 minutes, then were left to cool for 10 minutes. Sections were incubated in rabbit anti-phospho STAT5 antibody (dilution 1:1000, polyclonal rabbit anti-phospho-STAT5, Tyr 694, #9351, Cell signaling Technology, Beverly, MA, USA) for 48 hours. Following this, sections were incubated in biotinylated goat anti-rabbit IgG (dilution 1:300, BA-1000, Vector Laboratories, Inc., Burlingame, CA, USA) for 90 minutes and then incubated in Vector Elite avidin-biotin-HRP complex (dilution 1:100) for 60 minutes and labelling was visualized with nickel diaminobenzidine tetrahydrochloride using glucose oxidase to create a black, nuclear precipitate. For GFP immunohistochemistry sections were incubated in anti-GFP antibody (dilution 1:20 000, polyclonal rabbit-anti GFP, A6455, Life Technologies, Grand Island, NPY, USA) for 48 hours at 4C. Section were then treated as for pSTAT5 immunohistochemistry. Chromagen immunohistochemistry was examined using a light microscope at either 10x or 20x and numbers of positively stained cells were counted in either two sections (MPOA, PVN) or three sections (ARC, VMN) per mice and anatomically matched between mice.

### Statistical analysis

Statistical analyses were carried out with GraphPad Priam software. All group sizes refer to biological replications. P<0.05 was considered statistically significant and all data are presented as the mean ± SEM. Rout outlier test was used to determine any outliers, and outliers were removed from the data group. Student’s t-test was used to analyse daily RWA activity with different wheel types, distance travelled in EPM and OFT, virgin daily RWA, pSTAT5 positive cell number, virgin bodyweight, uterus weight, and fetus number. Paired Student’s t-test was used to analyse EE and ambulation in early pregnancy and total overall RWA, fine movement and time spend engaging in different behaviors following prolactin or vehicle treatment. Two-way ANOVA followed by Sidak’s multiple comparison test was used to analyse the change in RWA from virgin (four days prior to mating) to early pregnancy (days 1-3 of pregnancy). Repeated measures ANOVA followed by Sidak’s multiple comparisons test was used to analyse prolactin effects on RWA, ambulation, XY beam breaks, fine movement in virgin state. One-way ANOVA was used to analyse daily EE and ambulation during the different weeks of pregnancy. Mixed effects analysis followed by Sidak’s multiple comparisons test was used to analyse changes in daily RWA, body weight and food intake across pregnancy.Mann-Whitney tests was used to analyse estrous cycle data, pup number during early lactation and GFP positive cell number.

---

**Figure 4 Supplementary 1:**
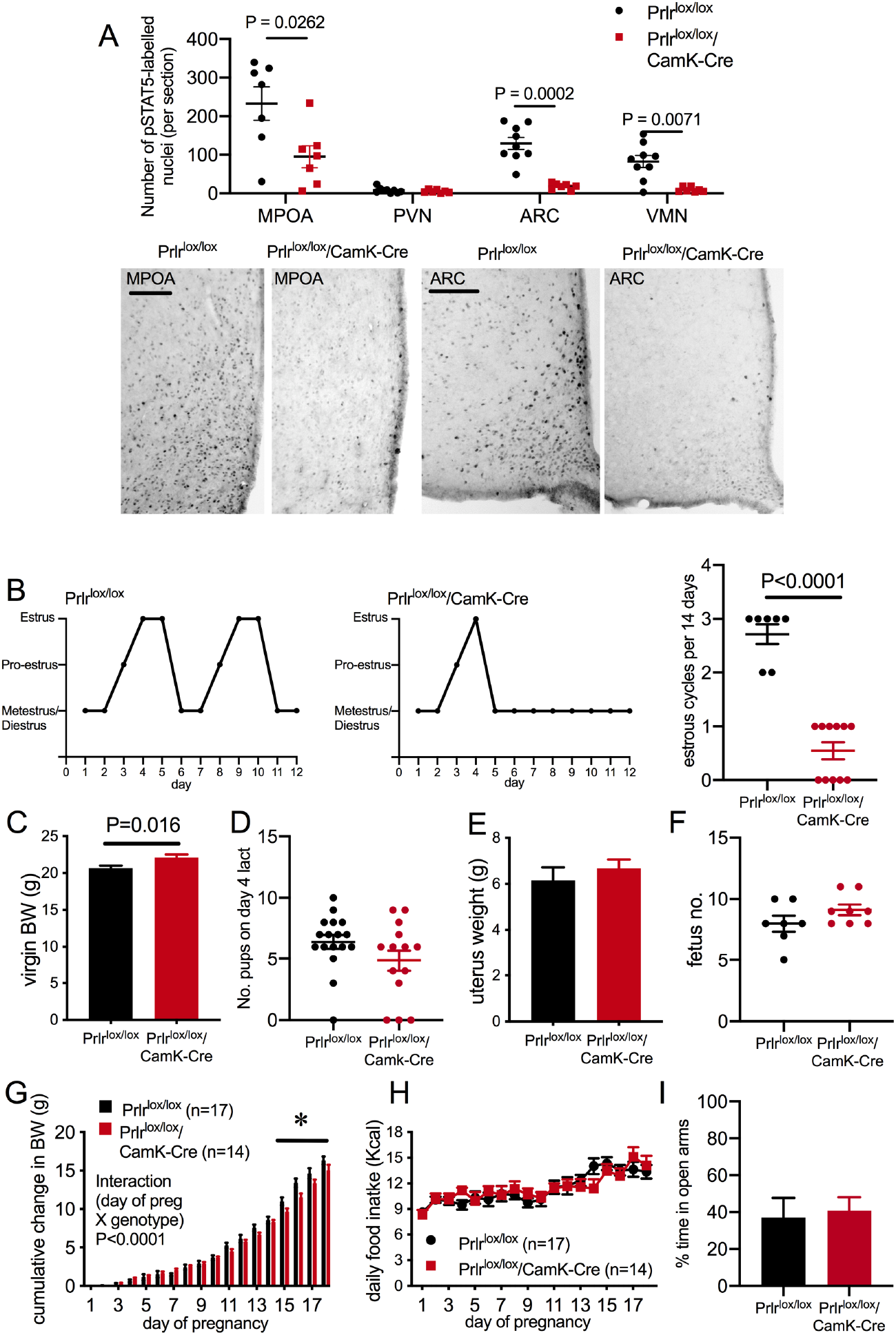
Conditional deletion of Prlr in forebrain neurons. **A** Bars represent numbers of prolactin-induced pSTAT5 (black nuclear staining) per section in various hypothalamic regions showing function loss of Prlr signaling in Prlr^lox/lox^/CamK-Cre female mice alongside representative images, n = 7-10 per group. **B:** Estrous cycles were disrupted in Prlr^lox/lox^/CamK-Cre female mice and showed long periods of continuous diestrus (n=7-11). **C**: Virgin Prlr^lox/lox^/CamK-Cre female mice (n=15) had a slightly but significantly larger body weight than control (Prlr^lox/lox^) female mice (n=17) of the same age. **D**: There was no significant difference in pup survived by day 4 of lactation in Prlr^lox/lox^/CamK-Cre and control mice. **E, F**: On days 15/16 of pregnancy Prlr^lox/lox^/CamK-Cre and control mice had similar uterus weights (**E**) and similar numbers of fetuses (**I**), n=7-8. **G, H**: While Prlr^lox/lox^/CamK-Cre mice showed a significant reduction in weight gain during pregnancy compared to control mice (**G**)(mixed model effect, significant interaction time x genotype P<0.0001, *post hoc* analysis to indicate specific times points where genoptyes are different using Sidak’s multipe comparison tests) there was no difference in food intake between the different genotypes (**H**) (two-way ANOVA). **I**: On an elevated plus maze in late pregnancy (days 15/16), Prlr^lox/lox^/CamK-Cre and control mice spent a similar amount of time in the open arms suggests no difference in anxiety-like behavior (n=7-8). Student’s t-test was used for A,C,E,F and I. Mann-Whitney test used for B and D.

**Figure 4 – Source data 1:**
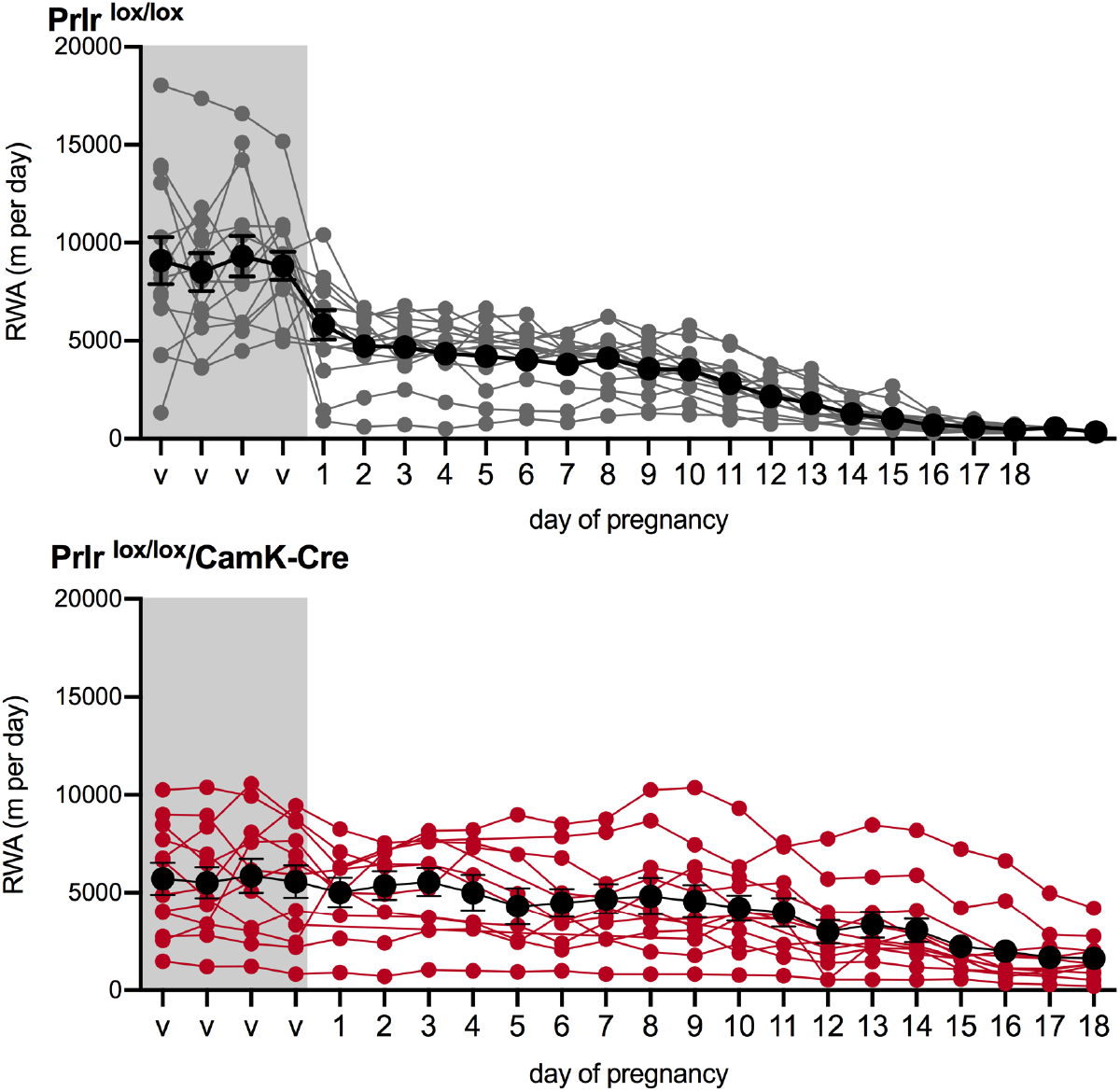
Each line (grey = control, red = Prlr^lox/lox^/CamK-Cre) represents total daily RWA from each individual mouse in the virgin state (4 days before mating) and during pregnancy (days 1-18). Black lines show the mean ± SEM for the group.

**Figure 4 – Source data 2:**
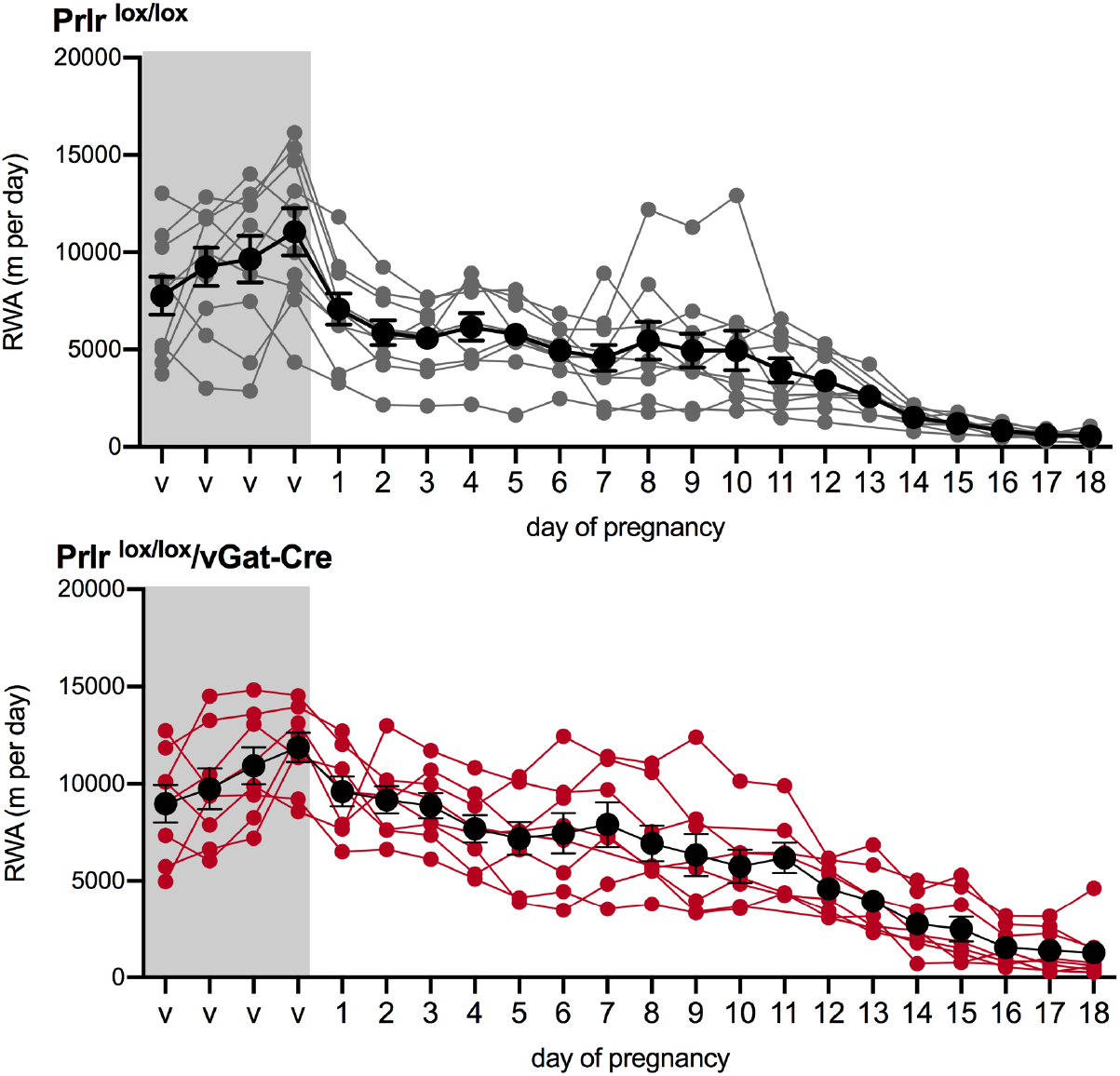
Each line (grey = control, red = Prlr^lox/lox^/vGat-Cre) represents total daily RWA from each individual mouse in the virgin state (4 days before mating) and during pregnancy (days 1-18). Black lines show the mean ± SEM for the group.

**Figure 4 Supplementary 2:**
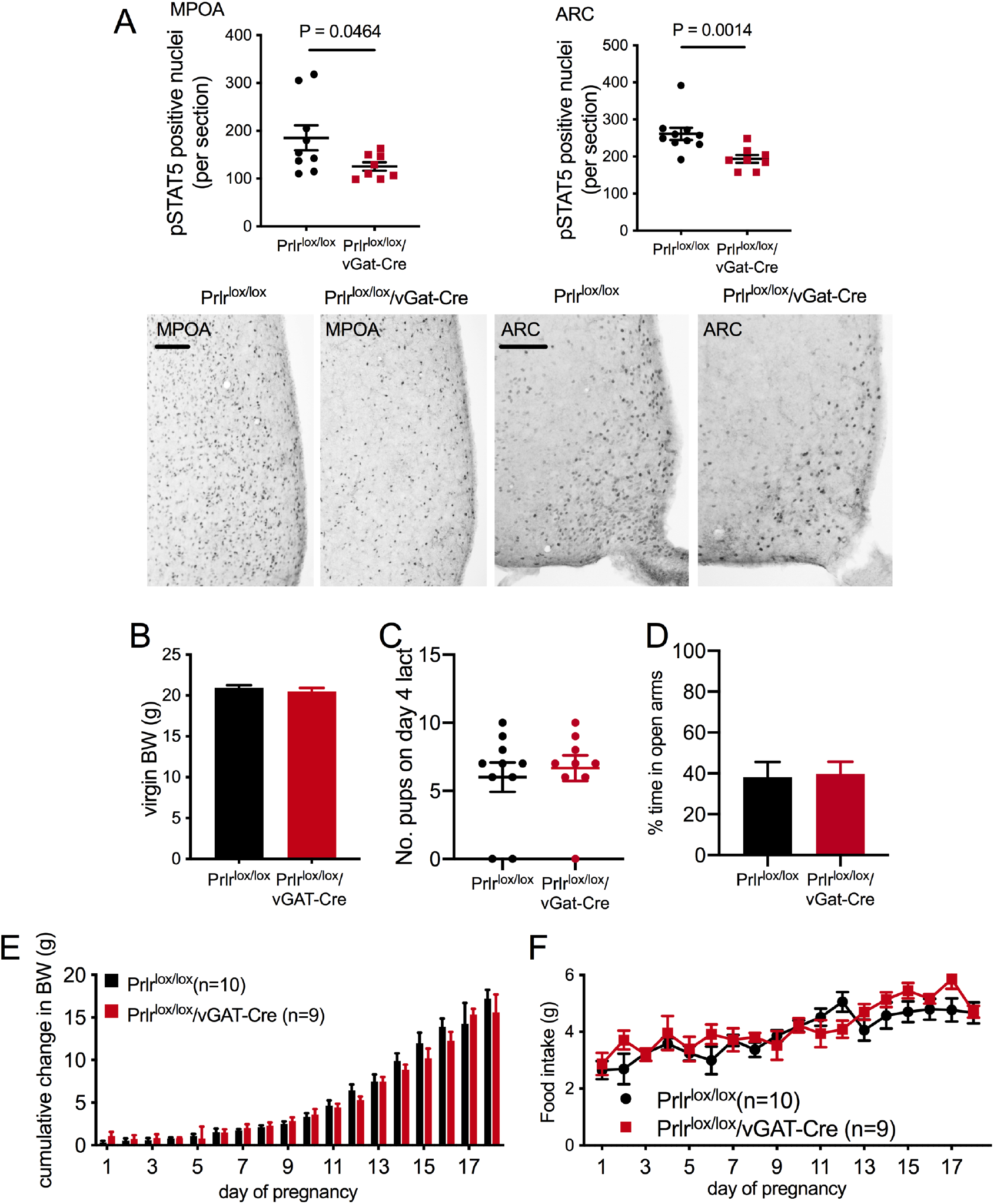
Conditional deletion of Prlr in GABA neurons. **A:** Bars represent numbers of prolactin-induced pSTAT5 (black nuclear staining) per section in hypothalamic regions showing function loss of Prlr signaling in Prlr^lox/lox^/vGat-Cre female mice alongside representative images, n = 8 per group (Student’s t-test). **B**: Virgin Prlr^lox/lox^/vGat-Cre female mice (n=8) and controls (n=10) had similar body weights. **J**: Following pregnancy and parturition both Prlr^lox/lox^/vGat-Cre and control mice had similar numbers of pups survive to day 4 of lactation (n=9-10). K: On an elevated plus maze in late pregnancy (days 15/16), Prlr^lox/lox^/vGat-Cre and control mice spent a similar amount of time in the open arms suggests no difference in anxiety-like behaviour (n=11-13). **L, M**: No differences in body weight gain (**L**) or food intake (**M**) during pregnancy were observed between the Prlr^lox/lox^/vGat-Cre (n=10) and control mice (n=9). A, B, D: Student’s t-test, C: Mann-Whitney test, E and F: mixed effects model.

**Figure 5 – Source data 1:**
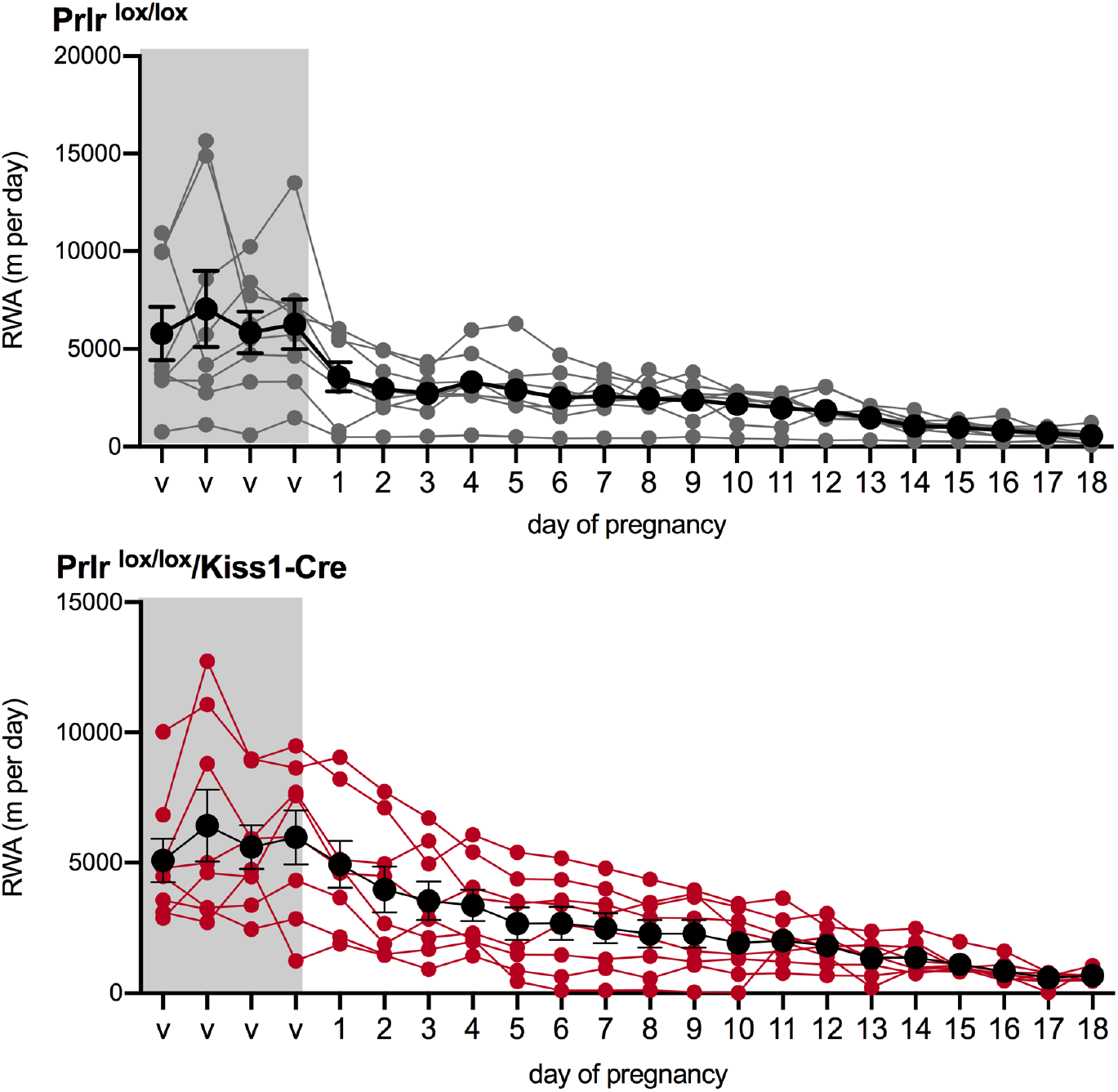
Each line (grey = control, red = Prlr^lox/lox^/Kiss1-Cre) represents total daily RWA from each individual mouse in the virgin state (4 days before mating) and during pregnancy (days 1-18). Black lines show the mean ± SEM for the group.

**Figure 6 – Source data 1:**
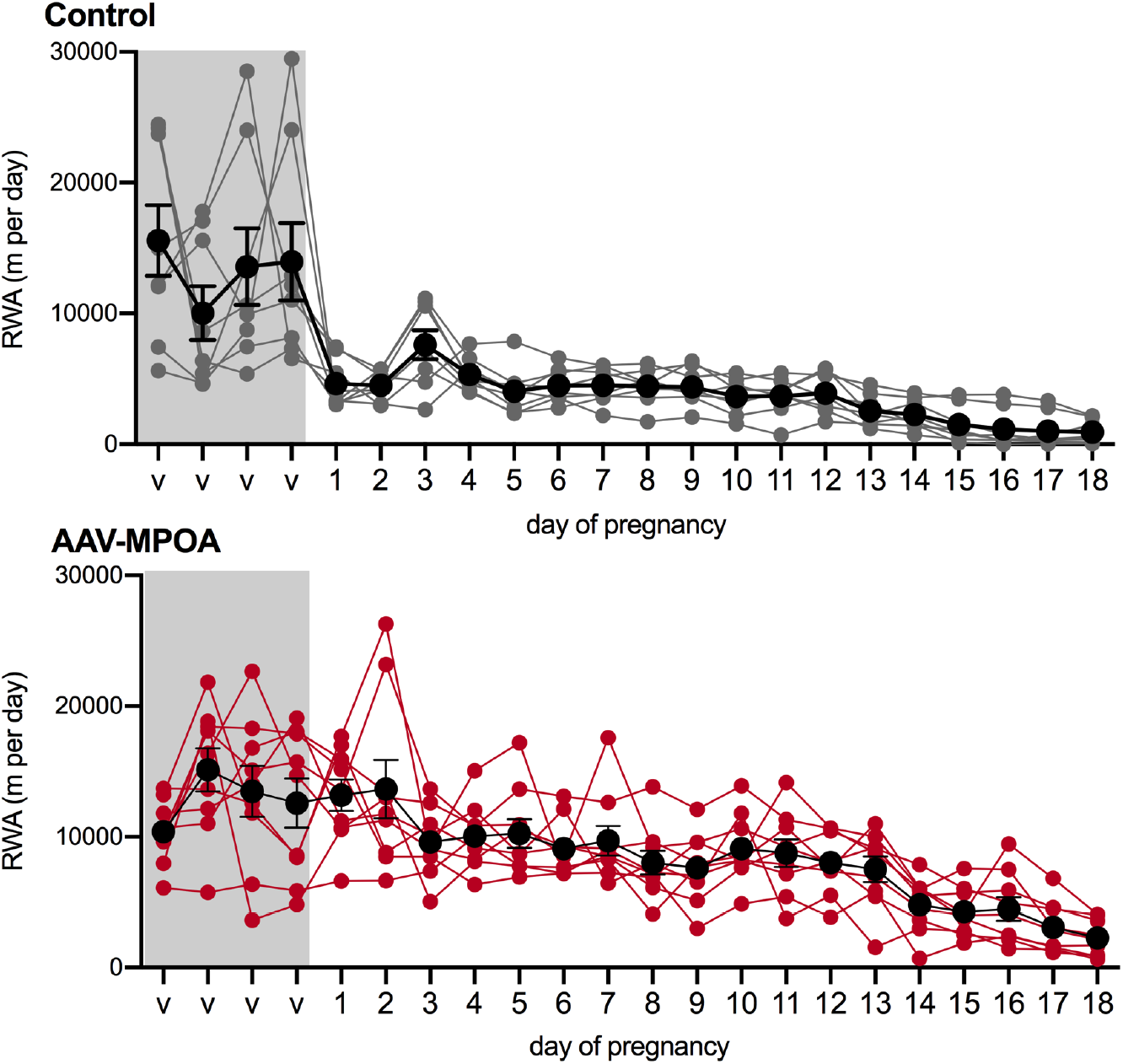
Each line (grey = control, red = AAV-MPOA injection of cre) represents total daily RWA from each individual mouse in the virgin state (4 days before mating) and during pregnancy (days 1-18). Black lines show the mean ± SEM for the group.

## Acknowledgments

We thank Pene Knowles for genotyping mice and Chantelle Murrell for assistance with immunohistochemistry quantification. This work was supported by a Health Research Council of New Zealand Programme Grant 14-568 (to DRG) and University of Otago School of Biomedical Sciences/Dunedin School of Medicine research grant funding (to SRL).

## Author contribution

**Sharon R. Ladyman**: Conceptualization, methodology, formal analysis, investigation, data curation, writing original draft preparation, supervision, project administration, funding acquisition

**Kirsten M. Carter**: investigation, formal analysis, data curation

**Zin Khant Aung**: investigation

**David R. Grattan**: Conceptualization, writing – review and editing, supervision, funding acquisition

